# Probing the DCAF12 interactions with MAGEA3 and CCT5 C-terminal degrons

**DOI:** 10.1101/2023.08.15.553295

**Authors:** Germanna Lima Righetto, Yanting Yin, Victoria Vu, David Duda, Magdalena Szewczyk, Hong Zeng, Yanjun Li, Peter Loppnau, Tony Mei, Yen-Yen Li, Alma Seitova, Jean-Francois Brazeau, Dalia Barsyte-Lovejoy, Vijayaratnam Santhakumar, Charu Chaudhry, Levon Halabelian

## Abstract

DCAF12 is the substrate recognition component of the CRL4 E3 ligase complex that can recognize C-terminal double-glutamic acid degrons to promote degradation of its cognate substrates via the ubiquitin proteasome system. MAGEA3 and CCT5 proteins were reported to be cellular targets of DCAF12. To further characterize the DCAF12 interactions with both MAGEA3 and CCT5, we developed a suite of biophysical and a proximity-based cellular NanoBRET assays showing that both MAGEA3 and CCT5 C-terminal degron peptides interact with DCAF12 in nanomolar affinity *in vitro and* in cells. Furthermore, we report here the 3.17 Å cryo-EM structure of DCAF12-DDB1-MAGEA3 complex revealing the key DCAF12 residues involved in C-terminal degron recognition and binding. Our study provides new tools and resources to enable the discovery of small molecule handles targeting the WDR domain of DCAF12 for future PROTAC design and development.

## INTRODUCTION

Protein degradation is a key housekeeping mechanism that allows proper cell function and homeostasis maintenance. The ubiquitin-dependent proteolysis system is essential for protein turnover, and relies on target protein ubiquitination as a prior step to proteasome-dependent degradation ^1^. The Cullin-RING ligases (CRL) are part of the largest family of ubiquitin ligases in eukaryotes, with extensive variability of protein members and degradation targets ^2,3^. The family shares a conserved architecture in which Cullin proteins – named from CUL1 to CUL7 - serve as a scaffold for E2 and E3 ligases, as well as RING finger and adaptor proteins ^2,3^. The Cullin E3 ligases are responsible for recruiting proteins for degradation, while the E2 ligases regulate ubiquitin transfer to the target protein.

DCAF12 (DDB1- and CUL4-associated factor 12) is a component of the CUL4 E3 ligase complex, which is known to have DDB1 (Damaged DNA binding protein-1) as the adaptor protein connecting Cullin 4, E3 ligase and target protein ^4^. Although DCAF12 biological function remains fairly understudied, recent reports have linked DCAF12 with neurotransmitter release ^5^ and spermatogenesis ^6^. DCAF12 is known to recognize double glutamic acid (-EE) C-terminal residues on substrate proteins, leading to their recruitment to the E3 ligase complex, ubiquitination, and subsequent degradation via the proteasome complex ^7,8^.

DCAF12 protein is a member of the WD40-repeat (WDR) family and shares the characteristic β-propeller/donut shaped fold ^9^. The WDR protein family has recently emerged as a promising target class for drug discovery, as several members of the family are implicated in key biological functions and have good draggability scores ^9,10^. In addition, E3 ligases and several other proteasome-related proteins have been in the spotlight for their potential to expand the druggable proteome by enabling Targeted Protein Degradation (TPD) ^11,12^. The TPD strategy involves hijacking the E3 ligase function by creating small molecule degraders – molecular glues or PROTACs (<u>Pro</u>teolysis <u>Ta</u>rgeting <u>C</u>himera) -that bind to the E3 ligase and recruit proteins of interest (POIs) for proteasome-dependent degradation ^1,13^. By combining the E3 ligase function and the druggable WDR domain, DCAF12 has the potential to be a successful target for drug discovery initiatives.

In light of recent findings elucidating the importance of C-terminal -EE degron for DCAF12 substrate recognition and binding for TPD ^7^, we decided to further characterize the DCAF12-substrate interaction both *in vitro* and in cell. Previous studies have shown that the presence of DCAF12 is directly linked to the decrease in MAGEA3 (Melanoma-associated Antigen 3) and CCT5 (T-complex Protein 1 Subunit Epsilon) protein levels in cells ^7^.

MAGEA3 is a member of a MAGE family of about 60 genes with conserved MAGE domain that are known to regulate RING-type E3 ligase activity by directly interacting with the RING ligase domain^14^. Even though MAGEA3 is normally expressed in male germ cells, the protein is also aberrantly expressed in various cancer types, including breast and lung cancer ^15^, multiple myeloma ^16^, and melanoma ^17 18^. Despite its implications in tumorigenesis^14^ and poor patience prognosis, the biochemical or structural characterization of MAGEA3 interaction with DCAF12 has not been reported.

Here, we show that DCAF12 can be successfully expressed and purified *in vitro* together with other members of the E3 ligase complex (DDB1 and DDA1) that are suitable for assay development and structural studies. We used a suite of biophysical assays, to characterize the binding for both MAGEA3 and CCT5 C-terminal degron peptides to DCAF12 WDR domain alone and in complex with DDB1 and DDA1. We also developed a cellular proximity-based assay to evaluate its target engagement in cells. Moreover, we generated the cryo-EM structure of DCAF12-DDB1-MAGEA3 complex revealing that the MAGEA3 C-terminal degron interacts with the conserved arginine and lysine residues located in the central cavity of DCAF12 WDR domain.

## RESULTS

### DCAF12 binds to MAGEA3 and CCT5 C-terminal degron with high affinity *in vitro*

DCAF12 is a member of the human CRL4 E3 ubiquitin ligase complex that is known to interact with the adaptor protein called DDB1 and acts as the substrate recognition component of the E3 ligase complex. To characterize the DCAF12 interaction with the reported substrates, we expressed the protein alone and in complex with DDB1 and other CRL4 E3 ligase family members. Despite our attempts to express and purify the full-length (FL) DCAF12 in isolation, we were only able to produce the N-terminally truncated form of DCAF12 (Δ1-78) in our hands (Figure S1A). Soluble expression of DCAF12-FL was achieved only in presence of DDB1, resulting in improved protein stability and purification yield (Figure S1A-B).

Previous studies have shown DDA1 as a core component of CRL4 E3 ligase complex by interacting with DDB1^19,20^ and making additional contacts with other members of DCAF substrate receptor proteins, as shown in DCAF15-DDB1-DDA1 structure^21–23^. To investigate the potential role of DDA1 in DCAF12-DDB1 complex stability, we co-expressed and co-purified his-tagged DCAF12-FL together with untagged DDB1-FL and DDA1-FL. The intact mass spectrometry analyses confirmed the presence of DDA1 together with DCAF12 and DDB1 after protein purification (Figure S1A-C).

In terms of the substrates, DCAF12 is known to promote ubiquitination and proteasomal degradation of MAGEA3 and CCT5 proteins in cells upon recognition of their C-terminal double glutamic acid degrons ^7,8^. To further characterize DCAF12 binding to MAGEA3 and CCT5 degrons, we developed a fluorescence polarization (FP)-based peptide binding and displacement assays *in vitro*. We designed small peptides harboring 11 C-terminal residues of MAGEA3 (LHEWVLREG<u>EE</u>) and nine from CCT5 (IRKPGES<u>EE</u>), both carrying the known -EE degron. In our FP direct binding assays, both MAGEA3 and CCT5 peptides showed potent binding to DCAF12 (Δ1-78), DCAF12-DDB1 and DCAF12-DDB1-DDA1 complex, with K_d_ values in the nanomolar range for both peptides (Figure 1A, Table S1).

**Figure 1:**
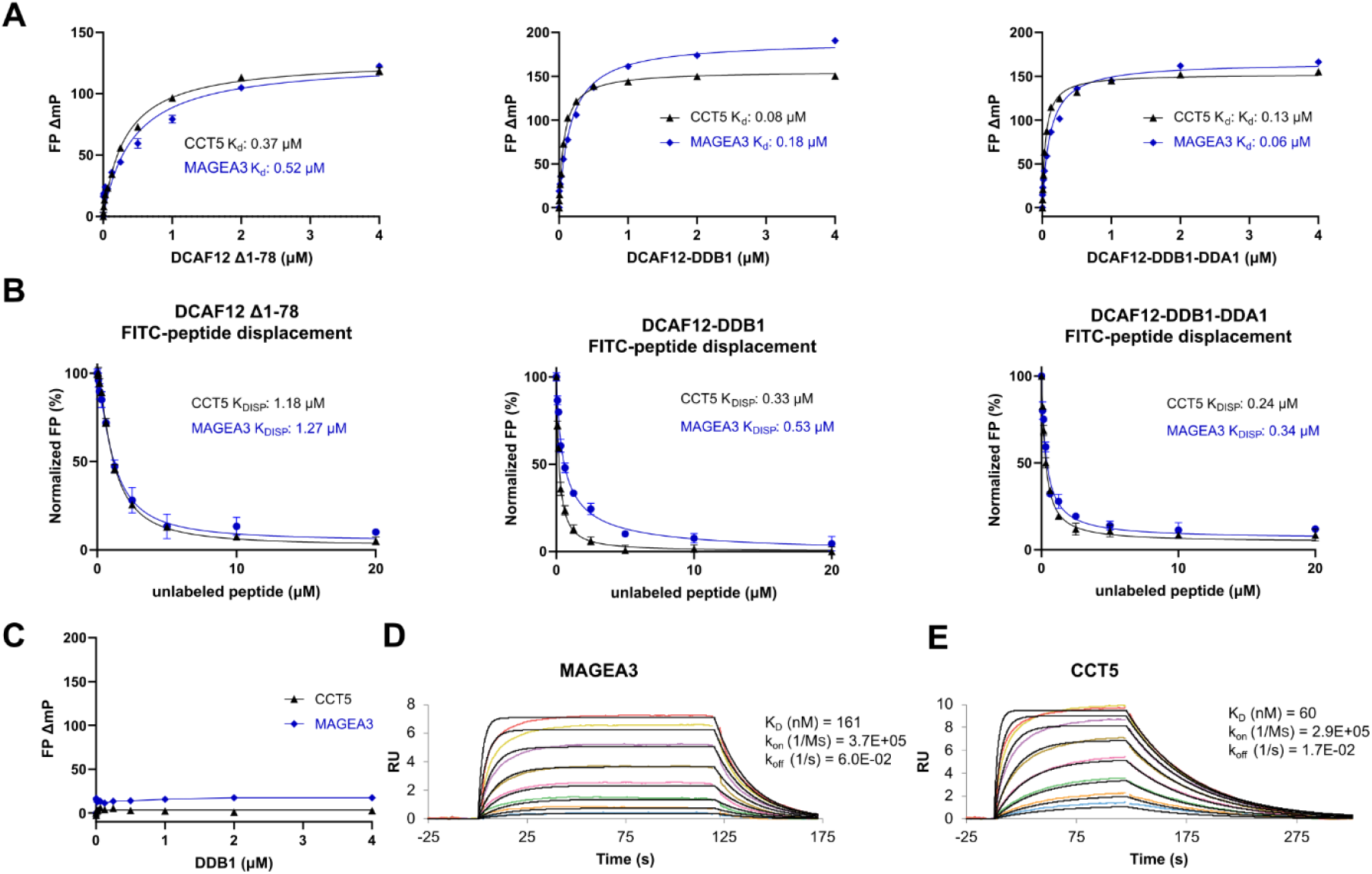
*In vitro* studies of DCAF12 interaction with MAGEA3 and CCT5 peptides. **A:** MAGEA3 and CCT5 peptides show similar binding affinities to DCAF12 alone and in complex with DDB1 and DDA1 in FP peptide binding assay. **B:** FP peptide displacement studies of DCAF12 bound to FITC-labeled MAGEA3 and CCT5 peptides with increasing concentrations of unlabeled peptides. **C:** CCT5 and MAGEA3 peptides show no binding to DDB1 when tested in FP assay. **D-E:** SPR analysis showing tight and similar binding of MAGEA3 and CCT5 peptides to biotinylated DCAF12 (Δ1-78).

Binding specificities were also evaluated by displacing the FITC-labeled peptides with the corresponding unlabeled peptides in a concentration dependent manner. Both FITC-labeled MAGEA3 and CCT5 peptides were successfully displaced in our assays by increasing concentrations of unlabeled peptides, confirming specific and reversible peptide-protein binding (Figure 1B, Table S1).

To rule out any direct involvement of DDB1 within the DCAF12-DDB1 complex in interaction with MAGEA3 and CCT5 peptides, we tested the direct binding activity of DDB1 alone in our FP assay using the same peptides. No interaction was observed between DDB1 alone and any of the tested peptides (Figure 1C), confirming that the peptides interact directly with DCAF12 only. In addition, we also tested the FP binding activity of DCAF12 to a control peptide, H3K9A (ARTKQTARASTG<u>GK</u>), which lacks the -EE residues at its C-terminal end. DCAF12 didn’t show any binding to the control peptide, thus ruling out any non-specific interaction between the FITC label and DCAF12 (Figure S2).

To further validate our FP assay results, we developed a surface plasmon resonance (SPR) assay using biotinylated DCAF12 (Δ1-78), in which both MAGEA3 and CCT5 peptides showed similar binding affinities comparable to the FP assay (Figure 1E).

### Conserved residues on DCAF12 central channel are essential for MAGEA3 binding in cells

Previous studies have shown that MAGEA3 and CCT5 expression is directly influenced by the presence of DCAF12, with DCAF12 knockout cells presenting higher expression for both proteins ^7^. In addition, the presence of two glutamic acid residues at the protein C-terminal end is key for MAGEA3 and CCT5 degradation, suggesting a direct role in protein:protein interaction with DCAF12 E3 ligase ^7^

To further investigate the DCAF12 interaction with MAGEA3 and CCT5, as well as to demonstrate cellular target engagement, we developed a cellular proximity-based assay using NanoBRET technology. To demonstrate the specific binding of MAGEA3/CCT5 to DCAF12 WDR domain, we explored suitable DCAF12 mutants. As no structure was available for DCAF12 at the time of our initial analysis, we took advantage of the predicted AlphaFold ^24,25^ structure. Considering the negative charge of the-EE side chains, our expectation was that the interaction between MAGEA3/CCT5 and DCAF12 would involve electrostatic interactions with positively charged, conserved residues within central pocket of DCAF12 WDR domain. A combination of electrostatic potential and sequence conservation analyses revealed that three arginine residues have key contributions to the highly positive surface charge of DCAF12: Arg203, Arg256, and Arg344 (Figure 2A, B).

**Figure 2:**
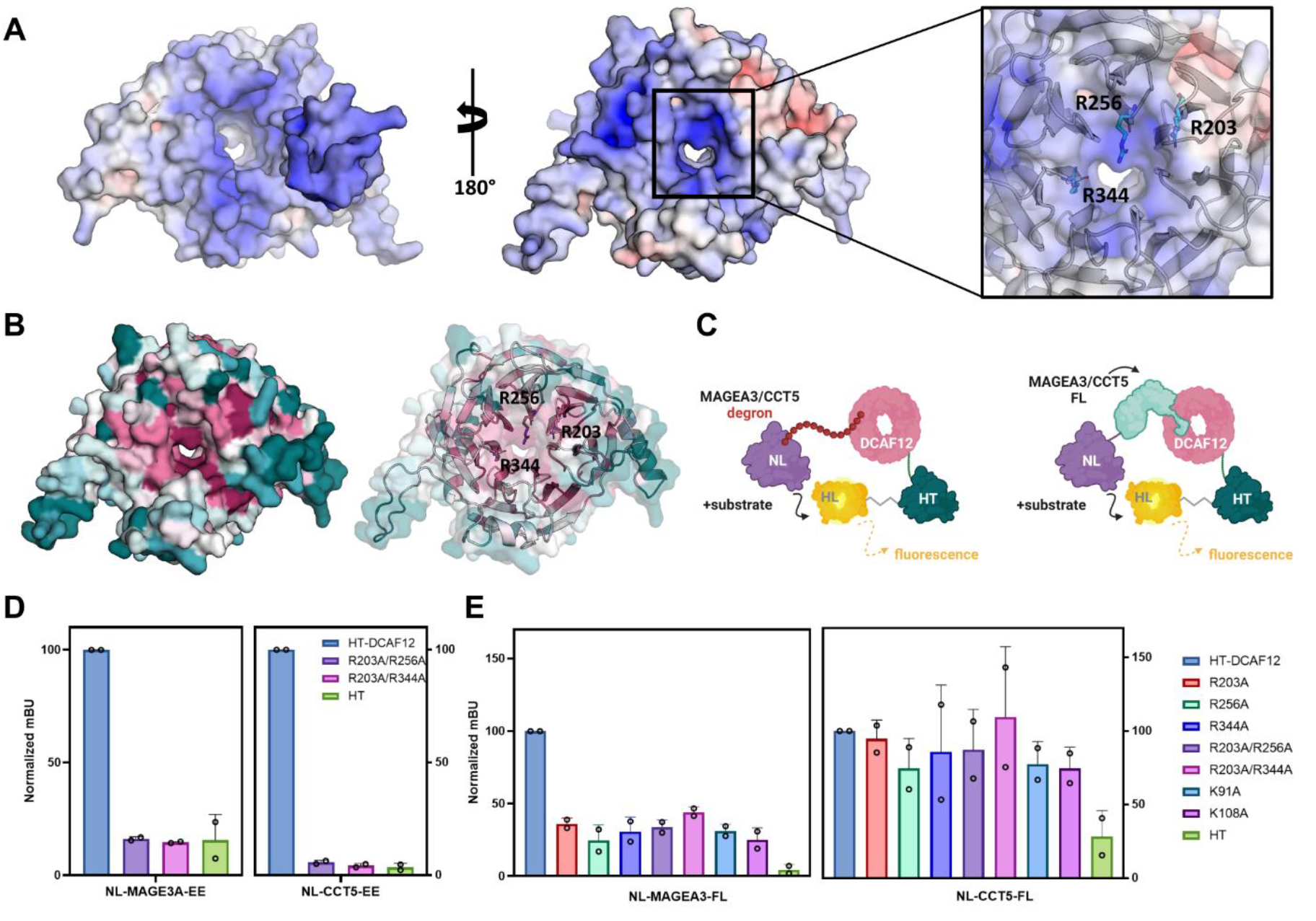
Conserved arginine and lysine residues near DCAF12 central channel participate in MAGEA3 binding in cells. **A:** DCAF12 electrostatic surface potential analysis showing negatively and positively charged residues colored in red and blue, respectively. On the left, DDB1 contacting surface of DCAF12. On the right, substrate contacting surface with positively charged residues –, Arg203, Arg256, and Arg344 -highlighted as sticks. Electrostatic potential is shown on a scale of −10 (red) kT/e to 10 (blue) **B:** Conservation analysis of DCAF12 showing high conservation of the same positively charged residues (represented as sticks) among homologous sequences. Residues were colored on a scale from variable/low conservation (dark teal) to high conservation (magenta). **C:** Schematic of NanoBRET strategy using MAGEA3/CCT5 FL protein or degron sequence cloned in frame with NanoLuc. DCAF12-WT and mutants were cloned in frame with HaloTag. **D:** NanoBRET assay using degron sequences of MAGEA3 and CCT5 against DCAF12 WT and combined arginine mutants R203A/R256A and R203A/R344A. **E:** Effect of DCAF12 mutations K91A, K108A, R203A, R256A, and R344A on the interaction with CCT5 and MAGEA3 FL proteins in NanoBRET assays in HEK293T cells.

We used site-directed mutagenesis to generate two DCAF12 mutants in the central pocket of the WDR domain: R203A/R256A, and R203A/R244A. DCAF12 wild type (WT) and mutants were tested in a NanoBRET proximity-based assay using both MAGEA3/CCT5 degron peptides (Figure 2C). Both DCAF12 R203A/R256A and R203A/R244A mutations impaired the interaction with MAGEA3 and CCT5 degrons, proteins, confirming the interaction is specific and the peptides bind to the central pocket of the WDR domain of DCAF12 (Figure 3D).

**Figure 3:**
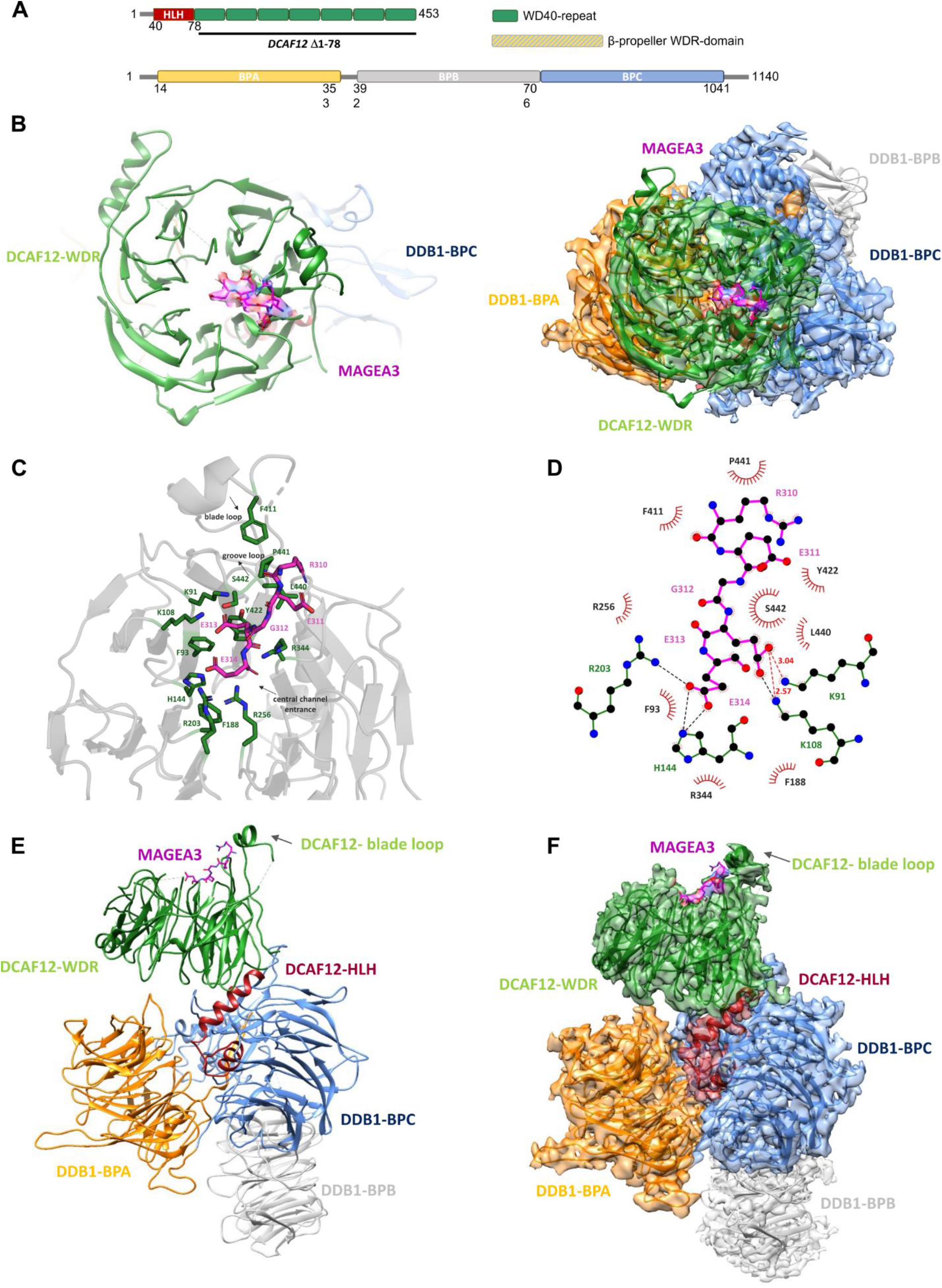
DCAF12-DDB1-MAGEA3 cryo-EM structure. **A:** Schematic of DCAF12 and DDB1 secondary structures with structural domains highlighted. **B:** Left panel, cryo-EM structure of DCAF12-FL bound to MAGEA3 peptide. DCAF12 is represented as cartoon and MAGEA3 as sticks. Right panel, frontal view of DCAF12-DDB1-MAGEA3 cryo-EM maps with model fitted. **C:** Cartoon view of DCAF12 (grey) bound to MAGEA3 C-terminal degron (magenta). Loops contributing to peptide binding are highlighted. DCAF12 residues contacting MAGEA3 peptide are represented as green sticks. **D:** Diagram of DCAF12 interactions with MAGEA3. MAGEA3 peptide residues are highlighted in magenta. DCAF12 residues making hydrogen bonds (red dashed lines) and salt bridges (black dashed lines) are colored green. DCAF12 residues involved in hydrophobic contacts are represented as red semicircles. Analysis was performed using LigPlot^+ 32^. **E:** Side view of DCAF12-DDB1-MAGEA3 cryo-EM model (left) and model fitted into maps (right).

To evaluate the ability of DCAF12 to interact with FL MAGEA3 and CCT5 proteins in cells, we repeated the assay using the FL proteins instead of the degrons peptides. In addition to the DCAF12 arginine mutants, the K91A and K108A mutants were also included in our assays, as both residues were shown to coordinate CCT5 binding to DCAF12 in a recently published DCAF12-DDB1-CCT5 cryo-EM structure ^26^. Interestingly, although the mutations decreased DCAF12 interaction with MAGEA3-FL, no changes were observed in its interaction with CCT5-FL (Figure 2E). The NanoBRET results suggest that positively charged residues located near the central channel of DCAF12 WDR domain are critical for the interaction with MAGEA3 and CCT5. Our NanoBRET assay using both MAGEA3/CCT5 degrons, as well as MAGEA3-FL, are suitable for determining cellular target engagement of the small molecules and will aid the development of cell active chemical probes.

### DCAF12 interaction with MAGEA3 relies on C-terminal -EE degron

To structurally characterize the DCAF12-DDB1 interaction with MAGEA3, we generated the 3.17 Å cryo-EM structure of the DCAF12-DDB1-MAGEA3 complex. Our structure confirms the seven-bladed β-propeller fold of DCAF12 (residues 79-453) that is characteristic of WD40 repeats domain ^9^ (Figure 3A, B). The interaction with MAGEA3 is driven by DCAF12 WDR domain, with the C-terminal degron residues of MAGEA3 interacting with DCAF12 central channel (Figure 3B). Interestingly, although MAGEA3 protein was co-expressed and co-purified as a 212 amino acids construct, only the last five C-terminal residues (REGEE) were observed in our cryo-EM map (Figure 3B-D). This result highlights the importance of the degron sequence for interaction with DCAF12.

A closer look into the DCAF12-MAGEA3 interaction site reveals that the C-terminal Glu313 and Glu314 residues of MAGEA3 pointing towards DCAF12 central channel, while the N-terminal portion points towards a small loop (residues 438-447) above the central channel (Figure 3C). The central channel of DCAF12 has two distinct patches of positively charged residues that play a crucial role in substrate recognition and binding. Notably, Arg256 and Arg344 residues enhance substrate binding by establishing interactions with the Glu314 backbone of MAGEA3 (Figure 3C, D). MAGEA3 Glu314 binding to DCAF12 is further stabilized by salt bridges with Arg203 and His144 residues (Figure 3D). On the other hand, Lys91 and Lys108 residues interact with Glu313 of MAGEA3 through a combination of hydrogen bonds and salt bridges (Figure 3C, D).

The N-terminal portion of MAGEA3 degron interacts with DCAF12 through several hydrophobic residues. MAGEA3 Arg310 and Arg311 participate in hydrophobic interactions with Leu440, Pro441 and Ser442 residues, forming a small loop above DCAF12 central channel (Figure 3C, D). MAGEA3 Arg310 is also involved in hydrophobic interactions with Phe411 residue, located in a loop (residues 401-414) formed between DCAF12 blades 6 and 7 (Figure 3C, D).

Beyond shedding light on DCAF12-MAGEA3 interaction, the cryo-EM structure also shows the conserved folding of DDB1, which is known to be formed by three β-propeller domains (BPA, BPB, and BPC) (Figure 3E, F). DDB1 BPA and BPC participate in the anchoring of DCAF12, which relies on a helix-loop-helix region (HLH) in DCAF12 N-terminus (Figure 3A, E). The DDB1-BPB β-propeller domain interacts with CUL4 scaffold protein in the E3 ligase complex and makes no contact with DCAF12 or other DCAFs (Figure 3E, F).

## DISCUSSION

E3 ligases are essential for cell homeostasis maintenance and protein turnover, as they function as key components of the ubiquitin/proteasome-dependent protein degradation system. Despite increasing interest in this target class for Targeted Protein Degradation, the great majority of E3 ligases remain underexplored for PROTAC development^27^. To enable the discovery of small molecule modulators/handles targeting the understudied DCAF12 E3 ligase, we sought to develop a toolkit by structurally and biochemically characterizing the interaction between the WDR domain of DCAF12 and its target degrons.

DCAF12 is a promising E3 ligase for TPD due to its ability to interact with various substrates by recognizing and binding to short peptide sequences (degrons) of target proteins, such as CCT5 ^7,26^, MAGEA3 ^7,8^, and the RNA helicase MOV10 ^6^. This interaction is mediated through the central pocket of DCAF12 WDR domain that could be targeted with small molecules handles for PROTAC design and development.

Previous studies have reported DCAF12-dependent degradation of MAGEA3 and CCT5 proteins in cells both using degron peptides and endogenous proteins ^7^. Our biophysical data demonstrates that DCAF12 interacts with MAGEA3 and CCT5 degron peptides with high affinity (Kd < 0.37 uM) *in vitro*. The similarity in binding affinities (< 5 fold difference) for DCAF12 alone, and in complex with DDB1/DDA1 suggests that DCAF12 WDR domain is directly responsible for degron binding, with no contribution from DDB1. Moreover, the full displacement of labeled degron peptides upon titration of unlabeled peptide confirms binding specificity and assay suitability as a toll for small molecule screening. A recent study reported CCT5 peptide binding to DCAF12-DDB1 complex using a TR-FRET assay with K_d_ and K_DISP_ values similar to our affinity determinations (< 3 fold difference) ^26^. To our knowledge, there are no current reports of biophysical characterization of MAGEA3 and DCAF12 interaction other than the one provided by our study.

Our NanoBRET assay results suggest that the WDR central channel of DCAF12 is responsible for interaction with the degron peptides, mediated by positively charged residues. The central pocket of other WDR proteins, such as DCAF1, WDR5 and EED, has previously been reported as highly druggable and implicated in protein:protein interactions ^9^. The nature of the DCAFs interaction with DDB1 through the N-terminal region occludes one side of the DCAFs central channel, leaving exposed for interactions only the upper side, as demonstrated by our cryo-EM data and previous DCAFs/DDB2-DDB1 structures ^21,28,29^.

Intriguingly, although mutations on DCAF12 central channel impaired the interaction with MAGEA3 degron and FL protein in our NanoBRET assay, only CCT5 degron responded to DCAF12 mutations. Our analysis shows that CCT5-FL has low engagement with DCAF12 in cells, with BRET signal 3.6-fold higher than the negative control, compared to 22-fold difference in signal from MAGEA3-FL. Moreover, DCAF12 mutations have little to no effect on their interaction, even when residues known to coordinate their binding were mutated ^26^. CCT5 protein belongs to a chaperonin family, and it is known to form an oligomeric complex containing two rings with 8 copies of the protein each ^30^. Interestingly, the CCT5 C-terminal degron residues from both rings seem to point towards the solvent-protected region of the oligomer (Figure S3), therefore making the -EE residues possibly inaccessible for interactions. Endogenous CCT5 protein levels were previously shown to increase upon DCAF12 knockout in A-375 cancer cell line ^7^, but the interaction could not be confirmed in our analysis using HEK293T cells. It is possible that only in certain circumstances the CCT5 degron is exposed for interaction with DCAF12. In line with this hypothesis, DCAF12-mediated MAGEA3 ubiquitination was observed to take place under starvation conditions ^8^.

Finally, our cryo-EM structure shows that DCAF12 forms a stable complex with MAGEA3, which was previously characterized as a DCAF12 degradation target ^7,8^. The structure highlights the importance of the C-terminal five residues of MAGEA3 for its recognition by DCAF12, with the last two glutamic acid residues (Glu313 and Glu314) playing a key role in its interaction with DCAF12. The positively charged residues Arg203, Arg256, Arg344, Lys91 and Lys108, situated at the central channel of DCAF12, contribute significantly to MAGEA3 binding by directly coordinating the terminal Glu313 and Glu314 residues. The importance of these residues was also demonstrated by our cellular NanoBRET assay, in which mutations in DCAF12 nearly abolished the interaction with MAGEA3 degron and the FL protein.

Our structural observations align well with a recently reported structure of DCAF12 in complex with CCT5 ^26^. The terminal glutamic acid residues of CCT5 (Glu541 and Glu540) also make direct contacts with Arg256, Lys91,and Lys108 residues of DCAF12^26^. Further, both MAGEA3 and CCT5 degron sequences share structural similarities in their binding to DCAF12, suggesting the existence of a conserved binding mechanism between the DCAF12 and its targets (Figure S4).

Our cryo-EM structure also highlights the conserved binding mode between DCAF12 and DDB1, with the N-terminal region of DCAF12 making interactions with DDB1-BPC and the WDR domain of DCAF12 interacting with DDB1-BPA and BPC. Similar binding mode was also observed for DCAF1 ^28^ and DCAF15 ^22^, as well as other DDB1 interactors ^31^. Although DDA1 was not included in our cryo-EM analysis, we expect similar interaction to DDB1-BPA, as previously observed in the DDB1-DDA1 ^19^ and DCAF15-DDB1-DDA1 ^23^ structures.

Overall, our study provides valuable tools for the development of small molecules targeting the WDR domain of DCAF12 for PROTAC development. First, we report the recombinant production of DCAF12 alone and in complex with interaction partners. Second, our FP and SPR assays will enable the screening and validation of compounds targeting DCAF12 in mid to high throughput format. Last, our NanoBRET assay using CCT5/MAGEA3 degrons and MAGEA3-FL will support target engagement evaluation of DCAF12 binders in cells through protein proximity.

## MATERIALS AND METHODS

### Protein cloning, expression, and purification

DCAF12 constructs were subcloned from Mammalian Gene Collection (MGC) stock BC063823 into baculovirus expression vector pFBOH-MHL. For co-expression with DCAF12, DDB1 (MGC stock BC063823) and DDA1 (MGC stock BC000615) were subcloned into pFBOH-LIC vector lacking any protein tags. For testing DDB1-FL direct binding to peptides on FP assay, the protein was expressed using pFBOH-MHL vector with his-tag. All protein constructs were expressed in *Spodoptera frugiperda* (Sf9) as previously described ^33^.

For protein purification, cells were resuspended in base purification buffer, supplemented with protease inhibitor, 0.1% NP-40, and benzonase (see Table S2 for buffer details). Suspended cells were lysed by sonication and clarified by centrifugation. Supernatant was collected and incubated with nickel or cobalt resin for 1 hour, under rotation at 4°C. Beads and supernatant were placed in an open column for affinity purification. Beads were washed with 10 to 20 column volumes (CVs) of base buffer, followed by a 5 CVs wash using base buffer supplemented with 5 mM imidazole. Protein was eluted using 5 CVs of base buffer supplemented with 250 mM imidazole. Protein was further purified by size exclusion chromatography (SEC) using base buffer. Purification samples were analyzed by SDS-PAGE. Clean fractions were pooled together, concentrated, and stored at – 80°C. Details about purification buffer and steps for each construct can be found on Table S2.

### Fluorescence Polarization assay

FP binding assays were performed using 20 mM Tris pH 7.5, 150 mM NaCl, 5% glycerol, 0.5 mM TCEP, 0.01% Triton buffer. All tested proteins were titrated from an initial 4 μM highest concentration, followed by a factor 2 serial dilution. FITC peptides (MAGEA3, CCT5, and H3K9A) binding to proteins was tested at 20 nM final concentration. Assay was incubated for 45 min at room temperature and plates were read using BioTek Synergy 4 (BioTek) plate reader. Prior to displacement assay, proteins at binding K_d_ concentration were mixed with FITC peptides at 20 nM final concentration. Displacement was evaluated by using a titration curve of unlabeled peptide, with 40 μM as highest concentration. Buffer, incubation time and reading conditions followed FP binding protocol. All FP analyses were performed in triplicates and, at least, two independent experiments. All K_d_ and K_DISP_ values were calculated using GraphPad Prism version 9.0.

### Surface Plasmon Resonance analysis

For SPR experiments, DCAF12 Δ1-78 sequence was subcloned into pFBD-BirA vector, expressed in *sf9* cells and purified as described (Table S2). Biotinylated DCAF12 (Δ1-78) was immobilized to a level of ∼3,200 RU on a Biacore Series S streptavidin chip (Cytiva BR100531) using a Biacore 8K+ (Cytiva). Experiments were performed in a running buffer of 10mM HEPES, pH 7.4, 150mM NaCl (Cytiva BR-1006-70), 3mM EDTA (Corning 46-034-CI), 0.005% P20 (Cytiva BR-1000-54), 5% Glycerol (Sigma G5516-1L) and 2% DMSO (Millipore Sigma MX1457-6). CCT5 and MAGEA3 peptides were injected over a period of 120s at 30 mL/min followed by a dissociation phase of 200s in an 8-point 2-fold dilution series with a top concentration of 1 mM. All steps were carried out at 15 °C. Reference and buffer-blank subtracted data were analyzed with the Biacore 8K Evaluation Software and fit to a 1:1 Langmuir model to obtain binding kinetics and K_D_ values.

### DCAF12 electrostatic surface and conservation analyses

DCAF12 electrostatic surface potential was generated using APBS ^34^ webserver. DCAF12 AlphaFold ^24^ predicted structure was prepared using ^35^ PARSE force field configuration. Automatic multigrid calculation was used to generate DCAF12 electrostatic potential. Residues conservation analyses were performed using ConSurf ^36^ server and DCAF12 AlphaFold predicted structure. Multisequence alignment was done using MAFFT method and homologs were selected from UNIREF90 using an identity cutoff of 35-95%. Conservation scores were calculated using Bayesian method. PyMOL (Schrödinger, LLC.) was used for further structure analysis and figure preparation.

### Nanobret assays using DCAF12 mutants and -EE degrons

HEK293T cells were plated in 96-well plates (2 × 104/well) and 4 h later transfected with

0.03 μg/well N-terminally HT-tagged DCAF12 (WT or mutants) or HT alone and 0.001 μg/well N-terminally NL-tagged CCT5 and MAGEA3 (FL or -EE degrons) using X-tremeGene HP transfection reagent (Roche), following manufacturer’s instructions. The next day, media was replaced with 40 μl of DMEM/F12 (no phenol red, supplemented with 4% FBS, penicillin 100 U/ml and streptomycin 100 μg/ml) +/-HaloTag® NanoBRET™ 618 Ligand (0.5 μl/ml, Promega). 4 h later, 10 μl of NanoBRET™ Nano-Glo Substrate solution (8 μl/ml Nano-Glo substrate, Promega, diluted in DMEM/F12 no phenol red, supplemented with 4% FBS, penicillin 100 U/ml and streptomycin 100 μg/ml) was added, and the signal was read. Donor emission at 450 nm (filter: 450 nm/BP 80nm) and acceptor emission at 618 nm (filter: 610nm/LP) was measured within 10 minutes of substrate addition using CLARIOstar microplate reader (Mandel). Mean corrected NanoBRET ratios were determined by subtracting the mean of 618/460 signal from cells without NanoBRET™ 618 Ligand x 1000 from the mean of 618/460 signal from cells with NanoBRET™ 618 Ligand x 1000. All DCAF12 mutants and -EE degron constructs were produced using Q5® site-directed mutagenesis kit (New England Biolabs)

### Sample preparation for electron microscopy

The QuantiFoil Au 1.2/1.3 300 mesh grids were glow discharged using PELCO easiGlow™ Discharge Cleaning System. A total of 3 μl of protein sample was applied onto the EM grids at a final concentration of 0.2 mg/ml. Sample vitrification was carried out using Vitrobot (Thermo Fisher Mark IV) with the following settings: blot time 3 s, blot force 3, wait time 0 s, inner chamber temperature 4 °C, and a 100% relative humidity. The EM grids were flash-frozen in liquid ethane cooled by liquid nitrogen. Cryo-EM data were automatically collected on a 200 kV Thermo Scientific™ Glacios™ microscope controlled by EPU software. Micrographs were captured at a scope magnification of 105,000X by a Facon4 detector (Gatan) operated in the counting mode. During a 6 s exposure time, a total of 40 frames were recorded with a total dose of 40 e-/Å2. The calibrated physical pixel size was 0.948 Å for all digital micrographs.

### Cryo-EM image processing and three-dimensional structure reconstruction

Cryo-EM data collection and image quality were monitored by the cryoSPARC Live v3.2 ^37^ installed in a local workstation. The image preprocessing steps included patch motion correction, patch CTF estimation, blob particle picking (50–150 Å diameter) and extraction, all performed simultaneously. A total number of 6777 raw micrographs were recorded during a 2-day data collection session using Glacios microscope. Acceptable 2D classes were further used as templates for particle repicking. Two rounds of 2-dimensional (2D) image classification were performed, resulting in ∼1.2 million good particle images (Figure S5A). Particles were used for 3D reconstruction. Four starting 3D models were calculated, with one major 3D class obtained. This major class was subjected to non-uniform 3D refinement and local refinement, with 476215 particles refined to a 3D map of 3.17 Å average resolution (Figure S5B). Detailed statistics about the cryo-EM experiments can be found on table S3 and Figure S5 C-E.

### Cryo-EM model building, refinement, and validation

Human DCAF12-DDB1-CCT5 cryo-EM structure (PDB 8AJN) was used as the initial model for atomic model building of the EM map. To create a starting model for the DCAF12-DDB1-MAGEA3 complex, each subunit of the previously solved protein complex was individually used for map fitting using UCSF Chimera ^38^. MAGEA3 degron peptide was modeled using CCT5 binding site as reference. Models were refined using PHENIX ^39^ and COOT ^40^.

## ACKNOWLEDGEMENTS

We would like to thank Ashley Hutchinson and Maria Kutera for their support with baculovirus protein expression. The Structural Genomics Consortium is a registered charity (no: 1097737) that receives funds from Bayer AG, Boehringer Ingelheim, Bristol Myers Squibb, Genentech, Genome Canada through Ontario Genomics Institute [OGI-196], EU/EFPIA/OICR/McGill/KTH/Diamond Innovative Medicines Initiative 2 Joint Undertaking [EUbOPEN grant 875510], Janssen, Merck KGaA (aka EMD in Canada and US), Pfizer and Takeda.

## COMPETING INTERESTS

Y.Y., D. D., J.F.B., and C.C., are or were employees of Janssen. Employees and past employees may hold stocks and shares or options to purchase them. All other authors declare no conflict of interest.

## Author Contributions

G.L.R. conducted protein construct design and purification, biophysical assay development, data analysis and interpretation, and wrote the manuscript. Cryo-EM experiments were performed, analyzed, and interpreted by Y.Y. and D.D. Site-directed mutagenesis and cellular experiments were designed and conducted by V.V. and M.S., with the assistance of T.M. H.Z. and Y-Y.L. assisted with protein purification. Y.L. and P.L. performed cloning for baculovirus production. A.S. coordinated baculovirus protein expression. D.B-L., V.S., J-F.B., C.C. and L.H. conceived the project, supervised the work and contributed to data interpretation. L.H., V.S., and C.C contributed to manuscript editing.

## SUPPLEMENTARY MATERIAL

**Figure S1:**
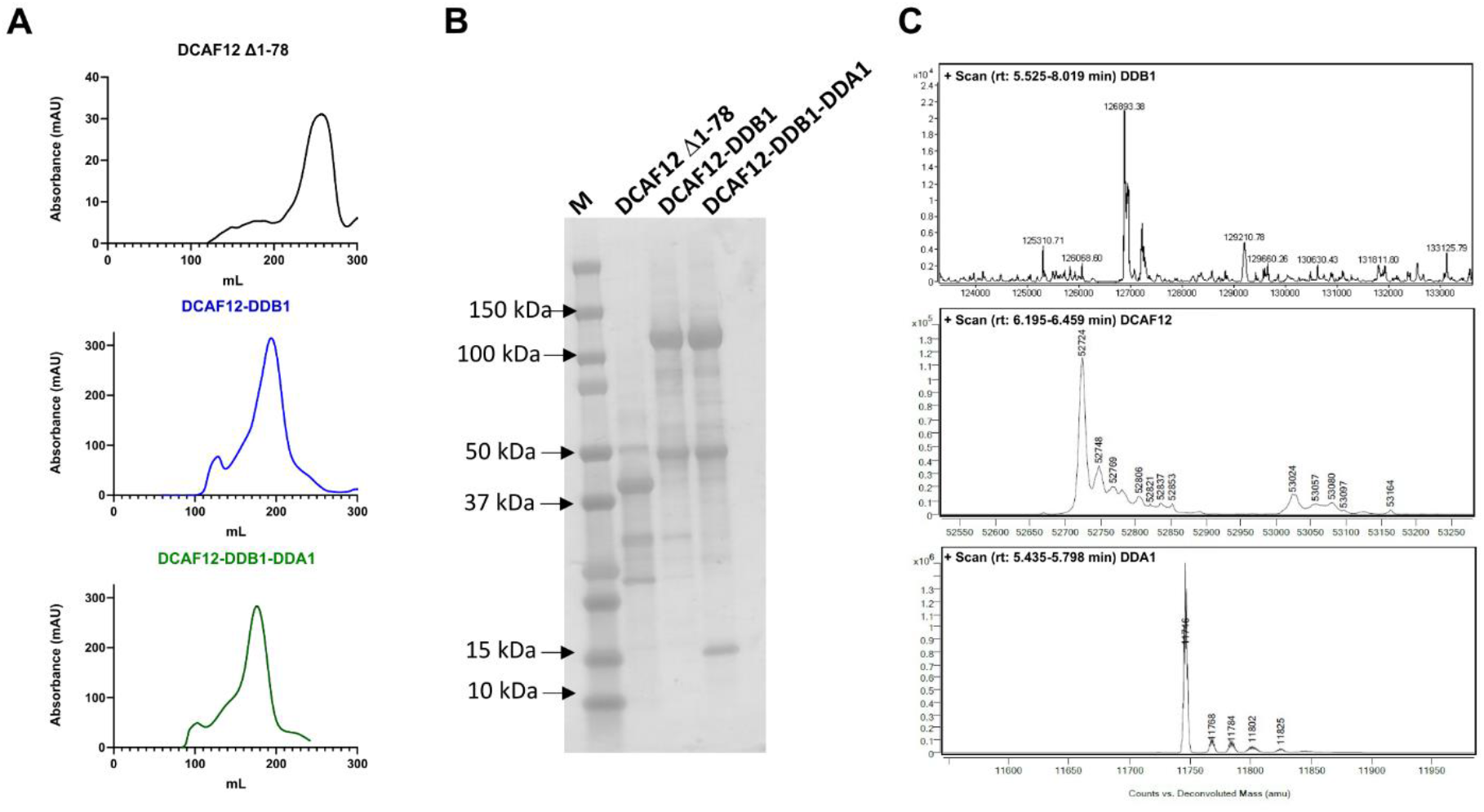
Generation of DCAF12 in complex with DDB1 and DDA1. **A:** Size exclusion purification profile of DCAF12 Δ1-78 (black), DCAF12-DDB1 (blue), DCAF12-DDB1-DDA1 (gray). **B:** SDS-PAGE analysis of final product of the size exclusion purifications shown in A. **C:** Intact mass LC-MS analysis of DCAF12-DDB1-DDA1 purification product showing peaks at expected molecular weight for each protein.

**Figure S2:**
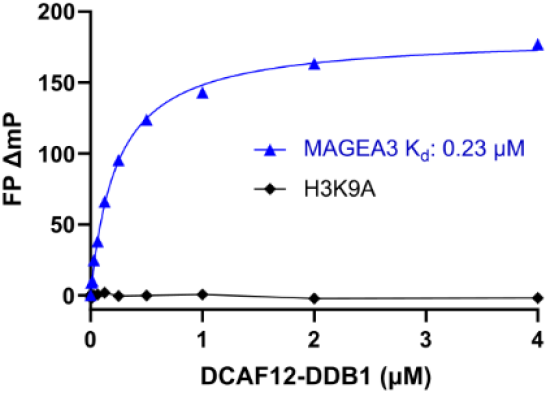
DCAF12-DDB1 binding to unrelated peptide lacking -EE degron. FP binding analysis comparing MAGEA3 and H3K9A binding to DCAF12-DDB1 complex.

**Figure S3:**
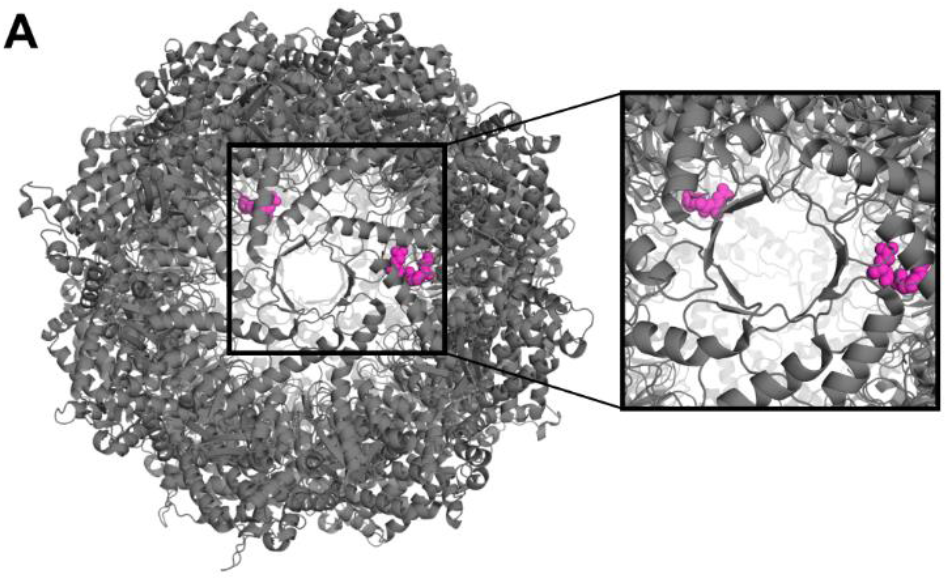
CCT5 C-terminal -EE degron is buried in oligomeric TRiC/CCT structure. Oligomeric CCT5 structure highlighting the position of C-terminal residues in the internal oligomer cavity (PDB ID: 7LUM). Terminal -EE residues are colored pink and shown as spheres.

**Figure S4:**
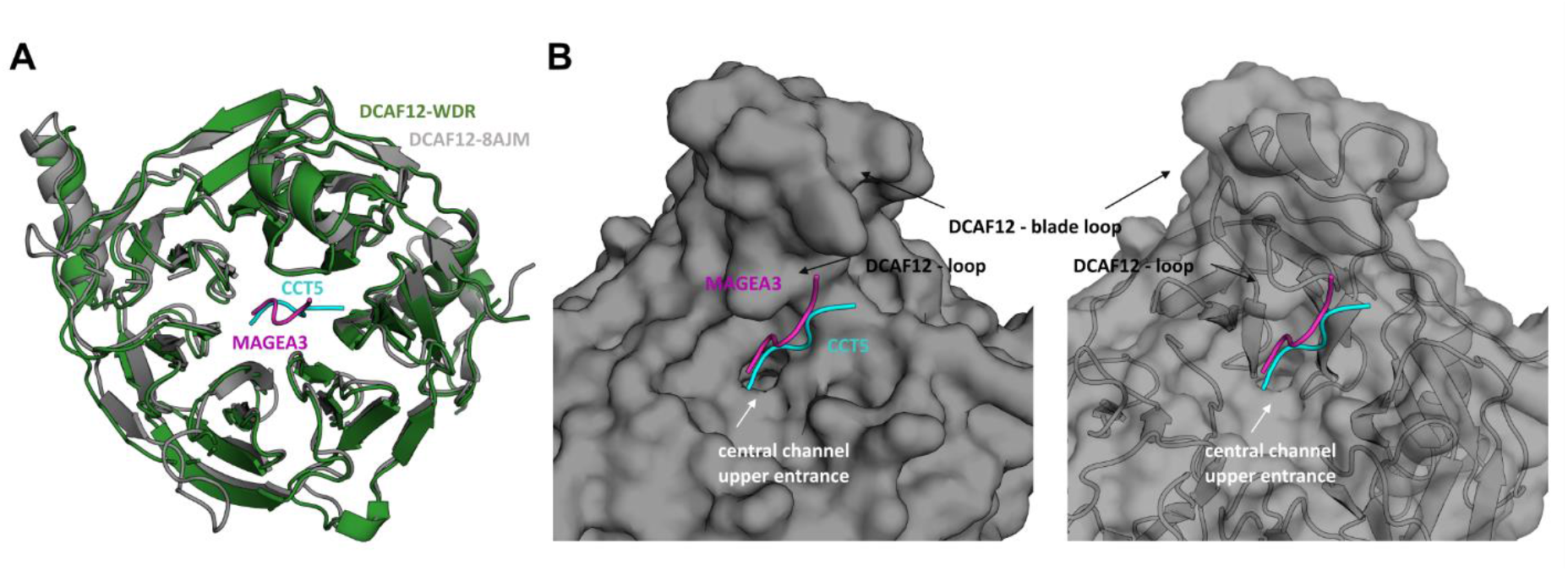
DCAF12 binding to MAGEA3 and CCT5 comparison. **A:** Structural alignment of DCAF12-MAGEA3 (green) and DCAF12-CCT5 structures (grey). Both MAGEA3 and CCT5 peptides bind DCAF12 in similar region, occupying part of the central channel. Alignment RMSD: 0.665 Å;. **B:** Close-up of MAGEA3 and CCT5 binding region on DCAF12 surface. Peptides can be seen contacting the top entrance of DCAF12 central channel. DCAF12 loops important to peptide binding are highlighted with arrows. MAGEA3 peptide is shown as cartoon in magenta, CCT5 is shown in cyan. DCAF12-DDB1-CCT5 PDB ID: 8AJM

**Figure S5:**
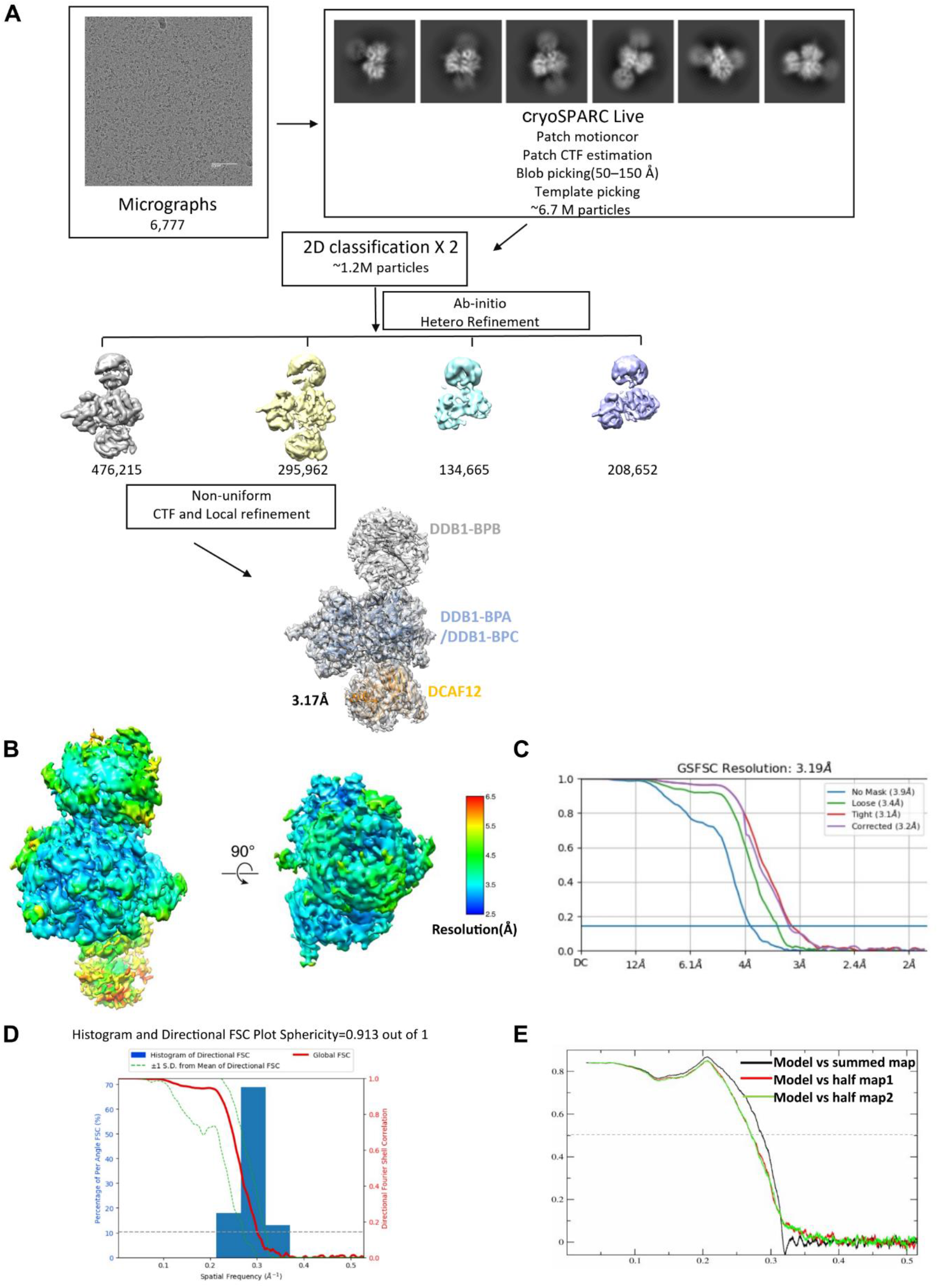
DCAF12-DDB1-MAGEA3 Cryo-EM data processing and refinement strategy. **A:** Workflow of image processing and 3D reconstruction. Representative micrograph from the DCAF12-DDB1-MAGEA3 collection is shown. Non-uniform analysis was applied to yield the 3.17-Å; resolution 3D map. **B:** Resolution estimations of the DCAF12-DDB1-MAGEA3 complex. Local map resolution was estimated using ResMap program and colored as indicated. **C:** The 0.143 criterion of the gold standard Fourier shell correlation (GSFSC) was used to estimate the average resolutions. **D:** Map directional anisotropy quantification using 3D-FSC server (https://3dfsc.salk.edu/). The sphericity of complex is 0.913, demonstrating good anisotropic property of the map. **E:** Model validation by FSC curves comparison. In red, comparison between model and half map 1 (work). Model and half map 2 (free) comparison is colored in green, and model and full map is plotted in black.

**Table S1:**
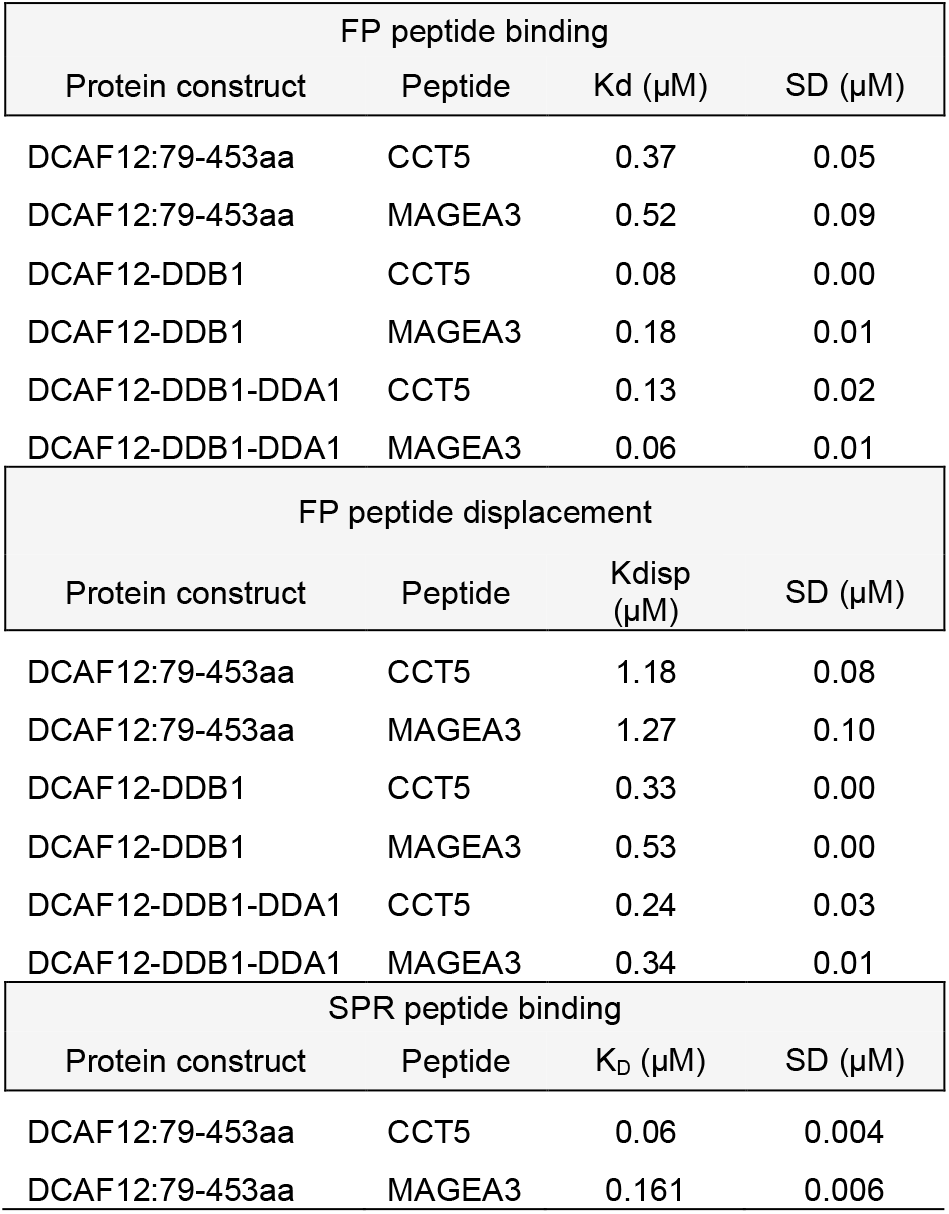
Biophysical data summary.

| FP peptide binding |  |  |  |
| --- | --- | --- | --- |
| Protein construct | Peptide | K<sub>d</sub> (μM) | SD (μM) |
| DCAF12:79-453aa | CCT5 | 0.37 | 0.05 |
| DCAF12:79-453aa | MAGEA3 | 0.52 | 0.09 |
| DCAF12-DDB1 | CCT5 | 0.08 | 0.00 |
| DCAF12-DDB1 | MAGEA3 | 0.18 | 0.01 |
| DCAF12-DDB1-DDA1 | CCT5 | 0.13 | 0.02 |
| DCAF12-DDB1-DDA1 | MAGEA3 | 0.06 | 0.01 |

| FP peptide displacement |  |  |  |
| --- | --- | --- | --- |
| Protein construct | Peptide | K<sub>disp</sub> (μM) | SD (μM) |
| DCAF12:79-453aa | CCT5 | 1.18 | 0.08 |
| DCAF12:79-453aa | MAGEA3 | 1.27 | 0.10 |
| DCAF12-DDB1 | CCT5 | 0.33 | 0.00 |
| DCAF12-DDB1 | MAGEA3 | 0.53 | 0.00 |
| DCAF12-DDB1-DDA1 | CCT5 | 0.24 | 0.03 |
| DCAF12-DDB1-DDA1 | MAGEA3 | 0.34 | 0.01 |

| SPR peptide binding |  |  |  |
| --- | --- | --- | --- |
| Protein construct | Peptide | K<sub>D</sub> (μM) | SD (μM) |
| DCAF12:79-453aa | CCT5 | 0.06 | 0.004 |
| DCAF12:79-453aa | MAGEA3 | 0.161 | 0.006 |

**Table S2:**
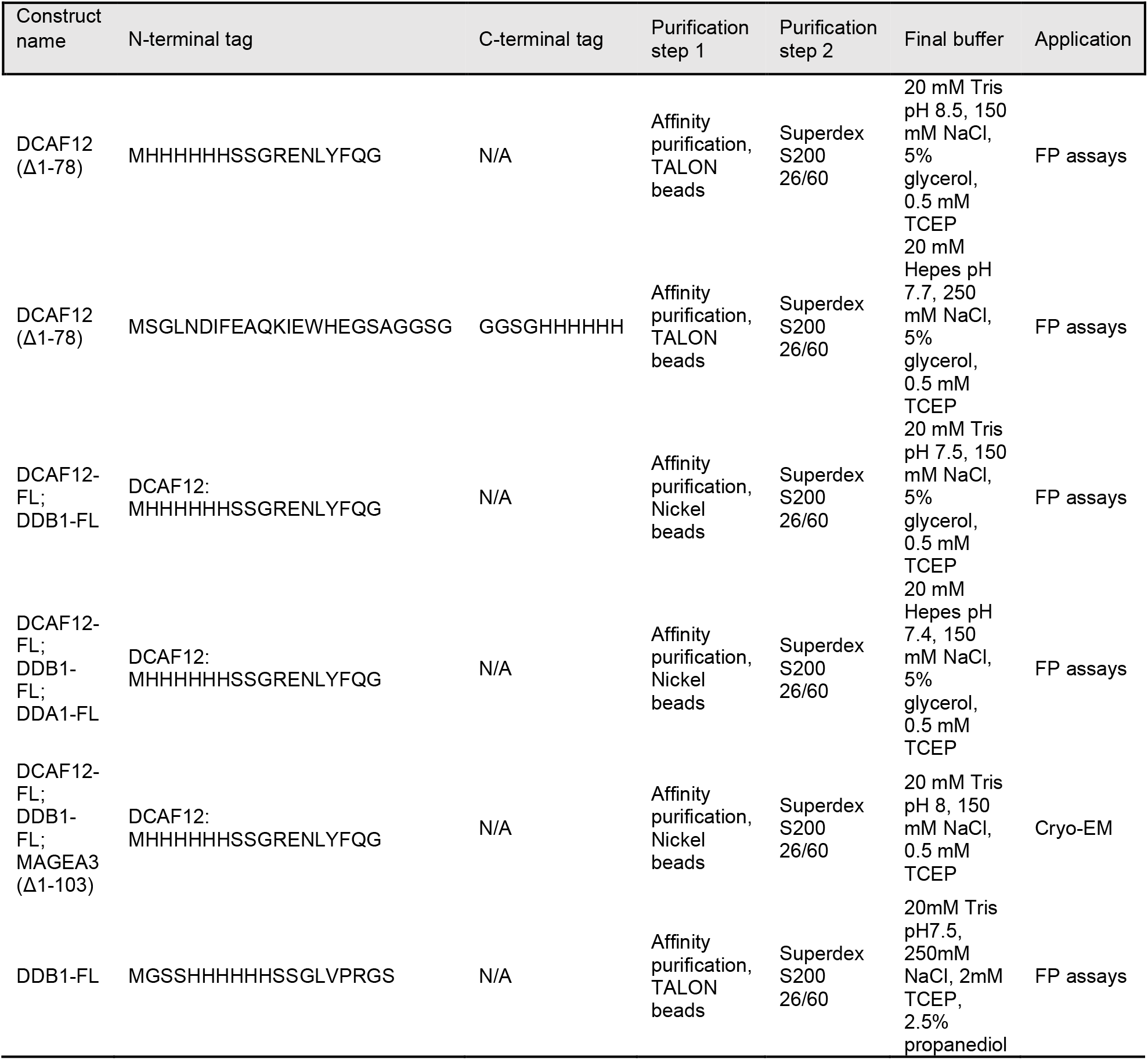
Protein purification information.

**Table S3:**
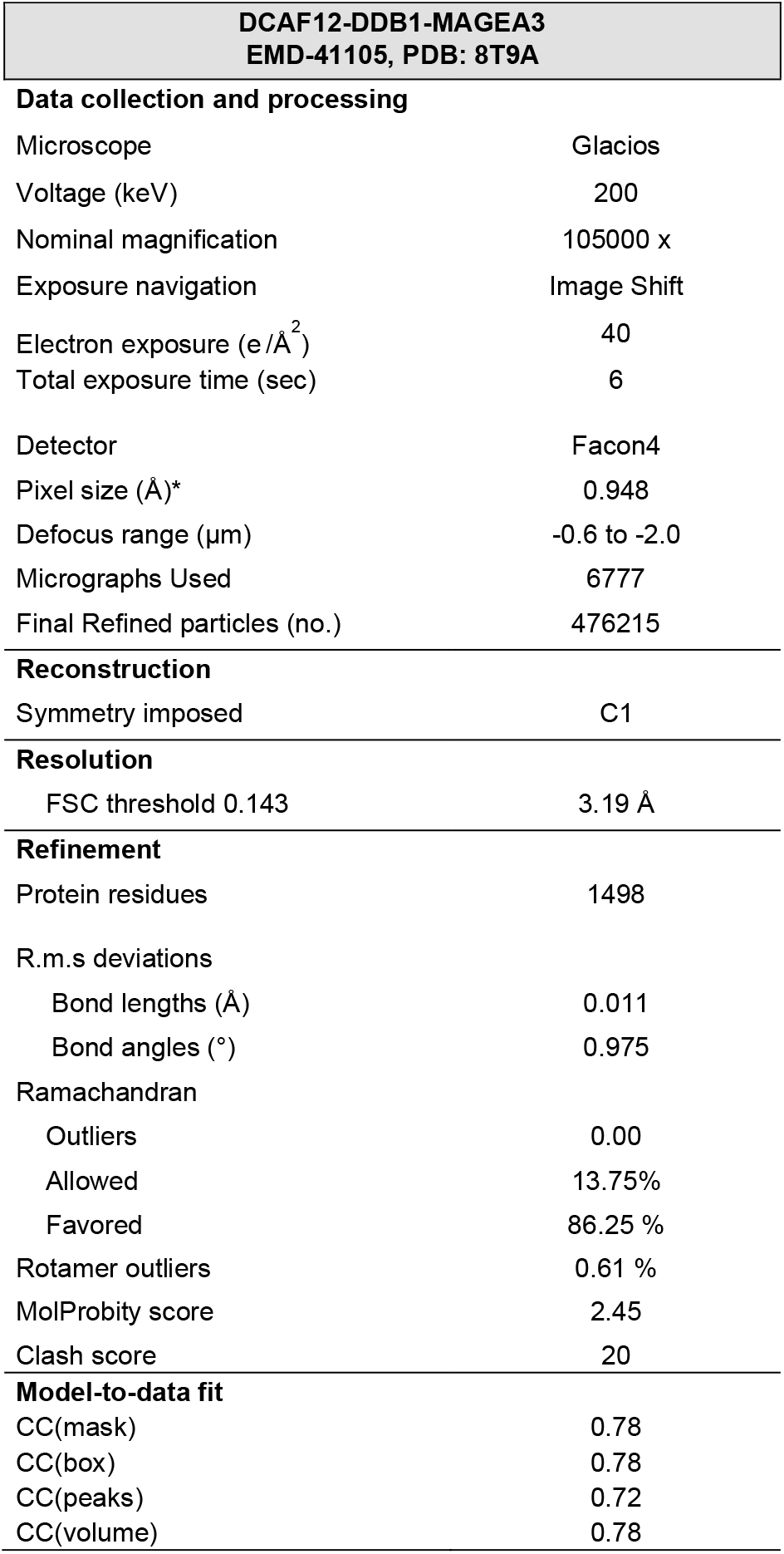
Data collection, reconstruction, model refinement, model-to-data fit statistics of Cryo-EM structures.

